# Probing sensitivity to statistical structure in rapid sound sequences using deviant detection tasks

**DOI:** 10.1101/2024.04.19.590221

**Authors:** Alice E. Milne, Maria Chait, Christopher M. Conway

## Abstract

Statistical structures and our ability to exploit them are a ubiquitous component of daily life. Yet, we still do not fully understand how we track these sophisticated statistics and the role they play in sensory processing. Predictive coding frameworks hypothesize that for stimuli that can be accurately anticipated based on prior experience, we rely more strongly on our internal model of the sensory world and are more “surprised” when that expectation is unmet. The current study used this phenomenon to probe listeners’ sensitivity to probabilistic structures generated using rapid 50 milli-second tone-pip sequences that precluded conscious prediction of upcoming stimuli. Over three experiments we measured listeners’ sensitivity and response time to deviants of a frequency outside the expected range. Predictable sequences were generated using either a triplet-based or network-style structure and deviant detection contrasted against the same set of tones but in a random, unpredictable order. All experiments found structured sequences enhanced deviant detection relative to random sequences. Additionally, Experiment 2 used three different instantiations of the community structure to demonstrate that the level of uncertainty in the structured sequences modulated deviant saliency. Finally, Experiment 3 placed the deviant within an established community or immediately after a transition between communities, where the perceptual boundary should generate momentary uncertainty. However, this manipulation did not impact performance. Together these results demonstrate that probabilistic contexts generated from statistical structures modulate the processing of an ongoing auditory signal, leading to an improved ability to detect unexpected deviant stimuli, consistent with the predictive coding framework.

## Introduction

Our brains are highly sensitive to predictable information. Detection of the regular patterns and structural arrangements in our environment is crucial for navigating daily life and forming long-term representations of the world. Learnable temporal structures allow us to make predictions about upcoming stimuli and anticipate sensory input before it arrives (Ekman et al., 2017), which improves neural and behavioral efficiency (Summerfield et al., 2008) and influences what enters conscious perception (Melloni et al., 2011). We extract, retain, and generalize consistent relationships between proximal and distal items in environmental input (Deocampo et al., 2019; Wilson et al., 2020), building concrete and abstract representations of the underlying structure (Daltrozzo & Conway, 2014; Dehaene et al., 2015). These “statistical learning” mechanisms are implicated in the organization and acquisition of language and other domains, such as action processing and object representation (Erickson & Thiessen, 2015; Romberg & Saffran, 2010; Sherman et al., 2020).

While it is known that humans and other organisms are sensitive to environmental statistical patterns, it is still unclear what type of statistics are learned under different situations, how these patterns are represented, and what impact the resulting representations have on ongoing sensory processing (Conway, 2020; Maheu et al., 2022; Planton et al., 2021). Research focused on auditory processing has historically emphasized deterministic sequences (i.e., precisely repeating frequency patterns; Barascud et al., 2016; Heilbron & Chait, 2018; Southwell et al., 2017; Southwell & Chait, 2018; Winkler & Denham, 2024), yet it is becoming increasingly clear that listeners track sophisticated statistics over multiple time scales (Benjamin et al., 2023, 2024; Maheu et al., 2019; Planton et al., 2021; Saffran et al., 1999; Skerritt-Davis & Elhilali, 2019). Thus, further research is needed to probe the impact of more complex temporal statistical relationships when processing rapid auditory sequences.

Sensitivity to structure is often probed behaviorally by using tokens that follow or violate previously encountered patterns in particular ways (Aslin & Newport, 2012; Conway, 2020; Frost et al., 2015; Milne et al., 2018; Sherman et al., 2020). This typically involves participants’ active engagement during the initial acquisition phase as well as subsequent introspection during the test period. An alternative approach is to embed deviant sounds in the input stream and measure the neural (Al Roumi et al., 2023; Daikhin & Ahissar, 2015; Furl et al., 2011; Garrido et al., 2013; Herrmann et al., 2015; Planton et al., 2021; Southwell & Chait, 2018; Tóth et al., 2023; Winkler et al., 1990) and/or behavioral responses during processing (Planton et al., 2021; Southwell & Chait, 2018).

For example, Southwell and Chait (2018) contrasted sequences of tones with no discernable pattern to deterministic sequences generated by iterating a 10-tone pattern. Identical deviants consisting of tones outside the main frequency range were embedded in both sequence types. For the predictable sequences, listeners were better and faster at detecting the deviant. Furthermore, application of EEG during passive listening showed an enhancement of the N1-P2-N2 deviant response suggesting that, despite the presentation of novel stimuli on each trial, the brain rapidly acquired the regularity and used it to facilitate deviant detection (also see Toth et al., 2023 for application in newborn infants). Similarly, Planton et al. (2021) generated increasingly complex binary sequences (i.e. containing only two different items) and found that detection of a novel tone frequency was impeded by sequence complexity. For both studies, deviants were more salient in sequences that could be more accurately predicted.

One hypothesis for this finding is that the response to deviants is driven by how confident the listener is in their prediction of ongoing sensory input (Yon & Frith, 2021). For contexts that can generate strong predictions, the listener will be more “surprised” than for contexts where they are more uncertain of the upcoming tones (Heilbron & Chait, 2018; Kanai et al., 2015). In the predictive coding framework this is referred to as “precision”, the reliability assigned to predicted sensory information (prediction error; Friston, 2005; Meyniel & Dehaene, 2017).

However, this previous work is limited by the use of relatively simple deterministic sequences and/or by administering sequences at a slow pace conducive to strategies that deliberately track the underlying structure. Consequently, it is difficult to establish if in the absence of conscious, intentional sequence tracking the automatic extraction of different forms of regularity are sufficient to form accurate predictions.

Here we probe how the incidental accumulation of complex statistical structures influences ongoing sensory processing. We tested if predictability would modulate frequency deviant responses in complex sequences under rapid presentation rates. Two types of probabilistic structures, a “Saffran” and “Community” structure were presented at a rapid rate (20Hz; figure 1c). This is substantially faster than the presentation rate usually employed in previous work (e.g. 4 Hz in Planton et al., 2021) and could be considered to surpass the threshold for conscious tracking of forthcoming stimuli (Warren et al., 1991) or at least prevent explicit prediction of upcoming tones, as can be done with stimuli such as syllables. Importantly, the deviant was a tone outside the main frequency range; therefore, its detection was not contingent on awareness of the structure. Moreover, employing such a task should direct attention to identifying extreme frequency changes, rather than seeking out patterns within the main tone stream, thus ensuring that any such learning of the patterns is incidental.

**Figure 1.**
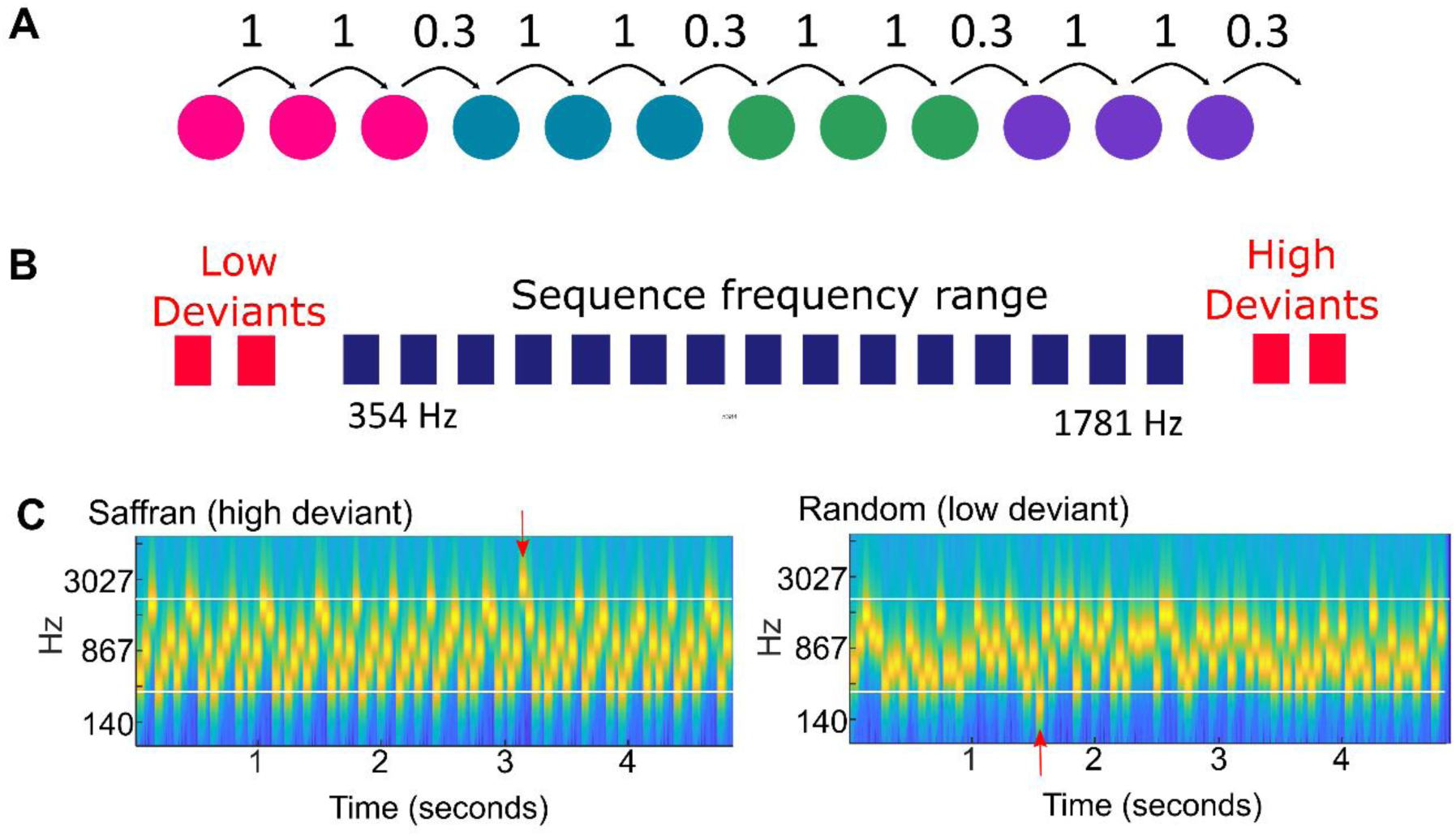
Stimulus design for Experiment 1 *Note.* Participants were presented with sequences of contiguous 50ms tone pips. The sequences were generated in two different ways: 1) predictable sequences generated according to the graph structure in **A** (example in **C**, left) and 2) randomly ordered sequences (example in **C**, right). To create the tone sequences, 12 tones were selected from a pool of 15 tones that were logarithmically spacing between 354Hz and 1781Hz (shown schematically in blue in **B**). The participants’ task was to detect an “outlying” tone, referred to as the “deviant,” which had a frequency outside the main frequency range (**C,** indicated by red arrows). In the deviant trials, one of four possible deviant tones was used (highlighted in red in **B**). ∼80% of trials contained a deviant.

The Saffran structure originates from the classic statistical learning paradigm, in which twelve items are organized into four triplets, which are then randomly ordered (Saffran et al., 1996). Consequently, tones occurring within a triplet are highly predictable relative to tones occurring at the boundaries between triplets. Neural evidence shows that the brain rapidly segments the stream using these triplets, even under passive conditions (Batterink & Paller, 2017). The second structure, a community structure, is more intricate (Benjamin et al., 2023; Karuza et al., 2017, 2019; Lynn et al., 2020; Schapiro et al., 2013). Community structures have recently attracted attention due to their closer resemblance to the statistics of natural environments (Girvan & Newman, 2002). In this community structure, each node is connected to an equal number of other nodes, but certain nodes are more interconnected, resulting in groups of nodes sharing contextual associations and being presented in close temporal proximity (Karuza et al., 2019; Schapiro et al., 2013). Critically, transitional probabilities at community boundaries are uniform. Thus, parsing sequences necessitates a model that includes the associations between both adjacent and non-adjacent preceding items, weighting those associations based on the distance between items (Benjamin et al., 2023; Lynn et al., 2020).

In a visual community structure learning paradigm, Karuza et al. (2017) investigated the impact of different traversal methods on learning. They implemented three distinct conditions: Unrestricted (called Random in their paper), Eulerian, and Hamiltonian. In the Unrestricted walk condition, sequences were generated by following any legal transitions, while in the Eulerian walk, every connection was traversed during each circuit. Both the Unrestricted and Eulerian walks result in nodes being revisited in close temporal proximity—a feature potentially crucial for detecting higher-order structure. In contrast, the Hamiltonian walk involved visiting each node exactly once during each circuit, leading to the absence of temporal cues signaling higher-order structure. These three traversal methods also resulted in sequences with varying levels of uncertainty. For the Eulerian and Hamiltonian paths, the likelihood of encountering a specific node or connection steadily increases if it has not already been encountered in the sequence. Consequently, as the possible routes through the network narrow systematically, there is a corresponding decrease in uncertainty (or increase in precision) regarding future stimuli. In contrast, for the Unrestricted sequences, there is no change in the number of viable options, and therefore uncertainty remains high.

The current study used both Saffran (Experiment 1) and Community (Experiments 2 & 3) structures. In contrast to deterministic sequences, the probabilistic nature of these types of structures should diminish overall precision, i.e., the certainty with which the subsequent tone can be predicted. In each experiment we also presented sequences that contained the same tones but in a Random order. Both types of structures should allow for more accurate predictions than the same set of tones presented in a random order, eliciting a heightened surprise response to unexpected stimuli, and resulting in enhanced and more rapid deviant tone detection. Furthermore, we predicted that, in line with the predictive coding framework (Friston, 2005), listeners would be more sensitive to deviants in contexts that can be better predicted; thus, deviants would be more “surprising” in the community structure conditions that were lower in uncertainty (i.e., Eulerian and Hamiltonian). Finally, we predicted that if listeners became sensitive to the higher order community structure, they would have weaker predictions at community boundaries and thus be less sensitive to deviants that occur after a change in community.

## Experiment 1

Experiment 1 featured sequences of contiguous 50ms tones adhering to the probabilistic Saffran triplet structure. Building on previous findings (Southwell & Chait, 2018), which demonstrated superior and quicker detection of deviant tones outside the main frequency range in deterministic sequences compared to randomly ordered tones, our primary objective was to examine how the presence of the probabilistic Saffran structure would impact the processing of incoming information, specifically an unexpected deviant tone.

For deterministic sequences that can be predicted with absolute certainty, evidence suggests a reliance on top-down expectations, minimization of prediction errors and a suppression of responses in lower-level cortical areas when the sensory input and prediction align (Sohoglu & Chait, 2016). Consequently, a deviation from these predictions should trigger a ’surprise’ response (Friston, 2005). In random sequences, the model’s accuracy is reduced, consequently these predictive mechanisms are relied upon less, resulting in a weaker surprise response and therefore lower accuracy in detecting deviant tones.

To extend this prediction to probabilistic sequences, participants were presented with sequences of tones that either adhered to the Saffran structure or were arranged in a random order. Participants’ task was to make a manual (key press) response to the appearance of a tone outside the frequency range. We hypothesized that, akin to the previously studied deterministic sequences, participants leveraging the underlying statistical structure to model upcoming stimuli would exhibit improved detection of deviant tones, resulting in enhanced performance and faster deviant detection for the probabilistic sequences relative to the random sequences.

### Method

#### Sample Size Selection

We calculated sample size using G*Power software (Faul et al., 2007) for a Wilcoxon signed-rank test, matched pairs with two tails. Based on the similarity of this design to Southwell and Chait (2018), we used their behavioral results to guide effect sizes. Effect sizes were not reported but means and standard deviations were available for the reaction time (Group 1 = 347; SD = 15; Group 2 mean = 387, SD = 25) and these were inputted to G*Power, computing an effect size *d_z_* =1.8 (correlation was assumed to be 0.5). Conducting an a priori power analysis, with this effect size and *α*= .05; β = *.8* indicated that a sample size of 5 should be sufficient to detect significant effects. However, given that we were using a more complex pattern that is less predictable than that used in Southwell and Chait (2018), as well as conducting the research online, we opted to recruit 20 participants to ensure the sample size would be sufficient.

#### Participants

Participants were recruited via the Prolific platform (prolific.com). One participant was excluded due to using the wrong operating system (see next section). Participants could also be excluded if they missed more than 50% of the control trials; none were excluded in the experiment. Control trials were intentionally designed to be easy, so a high failure rate on these trials indicated potential equipment issues or lack of attention. Participants who met these exclusion criteria were still compensated for their time spent completing the experiment. The final sample comprised of 19 participants (Age: *M = 27.0, SD* = 3.9; males = 13, females = 6).

Participants were remunerated at a rate exceeding the minimum specified by Prolific.com. To increase participant’s motivation (Bianco et al., 2021), additional bonus payments were rewarded based on their overall accuracy.

The experimental procedures were approved by the research ethics committee of University College London (ID: 14837/001), and each participant provided informed consent online prior to their involvement in the study.

#### Stimuli

The experiments were run and hosted remotely using the Gorilla online platform (www.gorilla.sc; Anwyl-Irvine et al., 2020). To ensure consistent participation conditions, restrictions were implemented in Gorilla, which included: 1) the use of a Chrome Browser on desktop or laptop computers (no phones or tablets allowed) and 2) a minimum connection speed of four Mbps. In addition, during data analysis, subjects were excluded if they used macOS computers, as these devices have been found to exhibit high variance and delays in recording reaction time (Anwyl-Irvine et al., 2020).

To minimize the influence of low-level cues, the stimulus generation procedure for each experiment was repeated five times, resulting in five different sets of stimuli that were randomly allocated to participants. MATLAB (The MathWorks) was used to generate the stimuli, which were then uploaded to Gorilla as .flac files.

The stimuli consisted of sequences composed of contiguous 50ms tone pips. There were 96 tones in each sequence, creating a trial length of 4800ms. A set of fifteen tones logarithmically spaced between 354 and 1781Hz was used to generate the sequences (frequencies: 354, 397, 445, 500, 561, 630, 707, 793, 890, 1000, 1122, 1259, 1414, 1587, and 1781Hz), a span of 2.33 octaves.

Three types of sequences were generated:

1. Structured sequences: These sequences were generated based on the Saffran triplet structure in Figure 1A and described in more detail below.
2. Random sequences: These sequences were created by taking a predictable sequence and randomly permuting the order of the tones. Each random sequence was paired with a corresponding predictable sequence.
3. Control sequences: These sequences consisted of 50ms tone pips of a single, randomly chosen tone frequency. The same tone frequency was repeated throughout the duration of the trial. This condition also served as an attention check, where any participant who missed more than 50% of control deviants in a given block was excluded from the analysis.

For each sequence type, deviant trials were generated. These deviant trials involved replacing one tone with a tone of a different frequency (deviant) falling outside the frequency range of the main sequence. The deviant could be one of four tone frequencies: two below (222Hz and 250Hz) and two above (2508Hz and 2809Hz) the main sequence range (Figure 1B and C) (0.5 octaves between the main frequency band and closest deviant). Deviant trials contained a single deviant that, across trials, was evenly distributed within a time window that excluded the first 750ms and the last 500ms of the trial. This design ensured sufficient time to establish the pattern, evenly spreading the deviants across the time window ensured that evidence accumulation prior to the deviant was consistent across conditions. In sum, structured vs random sequences varied in the arrangement of tones. Deviants were physically identical across conditions, but we hypothesized would be rendered easier or harder to detect based on the context in which they were embedded.

##### Structured sequence design

The structured sequences were based on the triplet paradigm introduced by Saffran et al. (1999) and adapted to 50-ms tones by Milne et al. (2021). For each stimulus set, twelve tones were randomly selected from the pool of fifteen and arranged into four tone “words”. This assignment of tones to words remained consistent throughout the entire experiment. For each trial, sequences were generated anew by randomly ordering the tone words, with the constraint that the same word did not occur twice in a row. Consequently, tone words always transitioned to a different tone word. This design resulted in a probabilistic structure where the transitional probability (TP; the probability that tone “a” will be followed by tone “b” calculated as the frequency of a to b/frequency of a) within tones of a word was 1, and the TP at word boundaries was 0.33. These structured sequences are referred to as the “Saffran” condition.

#### Procedure

Before participating in the experiments, all subjects underwent a headphone screening following the protocol established by Milne et al. (2021) and a volume calibration, adjusting their computer volume from a low to comfortable volume, using a stimulus similar to the main experiment.

*Deviant Detection Task:* Participants were instructed to actively listen for a tone that differed in pitch from the main sequence and to press the spacebar on their keyboard as quickly as possible when they detected the deviant tone. To ensure a clear understanding of the task, participants were provided with an example sequence that contained a high-pitched deviant tone. They had the opportunity to listen to this example multiple times. Following the high-pitched deviant example, participants were then presented with three practice sequences. Feedback was provided after each practice trial to reinforce correct responses. Subsequently, participants repeated the same process for the low-pitched deviant tone. The examples and practice sequences in this phase contained sequences of randomly ordered tones and thus did not possess any underlying structure. A final practice block contained all trial types: two trials with a high-pitched deviant, two trials with a low-pitched deviant, one trial with no deviant, and one trial with a control sequence containing a deviant (a single repeating tone frequency).

Following the practice, the test phase was presented as four blocks of 48 trials. Within each block, there were 21 Saffran sequences (16 of which contained a deviant tone, ∼76%) and 21 random sequences (16 with a deviant). Additionally, there were six control trials. Trials were presented in a random order using Gorilla’s trial randomization.

All trials started automatically and after each trial the participant received feedback regarding their performance in the form of a written message. If they responded before the target appeared, they received the message “Too Early”. If they missed the deviant, they received “Missed it-keep trying”. If they responded within 900ms of the target, they received “Correct!”. If they responded between 900 and 1500ms after the deviant, they received “Can you be faster?”. If they responded after 1500ms, they received “Sorry that one was too slow”. If they withheld a response on the non-deviant trials, they received “Correct there was no deviant!” Finally, if they made a false alarm, they received “’Incorrect, there was no deviant’.

At the end of each block, participants were provided with an overview of their performance. Following this, a one-minute break timer was initiated, encouraging participants to begin the next block as soon as the timer ended. Participants were informed that if they achieved an accuracy rate of over 70%, they would receive a £1 bonus, and if they achieved an accuracy rate of 80%, they would receive a £2 bonus.

#### Data analysis

The statistical analysis focused on two main measures: sensitivity scores (d’) and reaction time. Sensitivity scores (d’) were computed using the hit and false alarm rates [z(hits) – z (false alarms)]. A keypress was considered a hit if it occurred between 100ms and 1500ms following a target gap. In cases where hit rates or false alarms reached ceiling values (1 and 0, respectively, resulting in an undefined d’), a standard correction was applied by adding or subtracting ½t (where t is the number of trials). Reaction times were recorded if they fell between 100ms and 1500ms post-deviant onset. To reduce the effect of individual differences in response speeds, reaction times were adjusted by subtracting the average RT to the control condition, for each block and each subject, only adjusted RTs are plotted. Both measures were calculated on a block-by-block basis and averaged across blocks before comparing across conditions. Since there were 16 “target” trials i.e. with deviant and 5 “no target” trials, i.e. without a deviant, the ceiling for d’ was 3.144.

Each measure underwent a pairwise statistical test to compare deviant detection in Saffran relative to Random conditions. Due to the non-normal distribution of sensitivity scores, a Wilcoxon signed-rank test was employed. Adjusted reaction times, being normally distributed, were tested using a paired samples t-test.

To investigate the emergence of these effects with increased exposure to the structure, a repeated-measures analysis of variance (RM-ANOVA) was conducted on d’ and reaction times (RT) across blocks. The factors included sequence type (Saffran, Random) and block number (1, 2, 3). Bonferroni corrections were applied to all pairwise tests.

#### Transparency and Openness

For all experiments we report how we determined our sample size, all data exclusions, all manipulations, and all measures in the study and JARS (Appelbaum et al., 2018). All data, analysis code are available at https://osf.io/f2td6 and research materials are available at https://app.gorilla.sc/openmaterials/824494 for Experiment 1 and https://app.gorilla.sc/openmaterials/824480 for Experiments 2 and 3. Data were analyzed using Matlab, version 2021b and SPSS, version 27. This study’s design and its analysis were not pre-registered. This data is likely generalizable to all populations that regularly use computers and have access to the internet as the data was collected online via prolific.com with no geographic restrictions, which is available to participants from all OECD countries, except Turkey, Lithuania, Colombia, and Costa Rica; it is also available in Croatia and South Africa. The demographic data provided by prolific indicated that participants were not limited to student populations; specific numbers are not reported here due to a larger number of missing cases for this demographic preventing accurate reporting. Data collection took place between 2020 and 2023.

### Results

There were higher levels of sensitivity (d’: *Z* = -2.951, p = .003, *d_z_* = -1.87) and faster reaction times (*t*(18) = -2.286, p = .035, *d* = -0.524) when detecting the deviant in Saffran compared to random sequences (Figure 2A, B). Introducing a factor of block found that regardless of sequence type, d’ improved over time (Main effect of block: *F*(3,54) = 7.961, p = .007, η_p_^2^ = .307, figure 2C). The effect of sequence type was retained (*F*(1,18) = 12.207, *p* = .003, η_p_^2^ = .404) with no significant interaction (*F*(3,54) = .2.825, *p* = 0.85, *η_p_^2^* = .136). While there was no significant interaction with block, visually it appears this effect is not present until after the first block (figure. 2D). Note some participants were at ceiling (d’ ceiling = 3.14).

**Figure 2.**
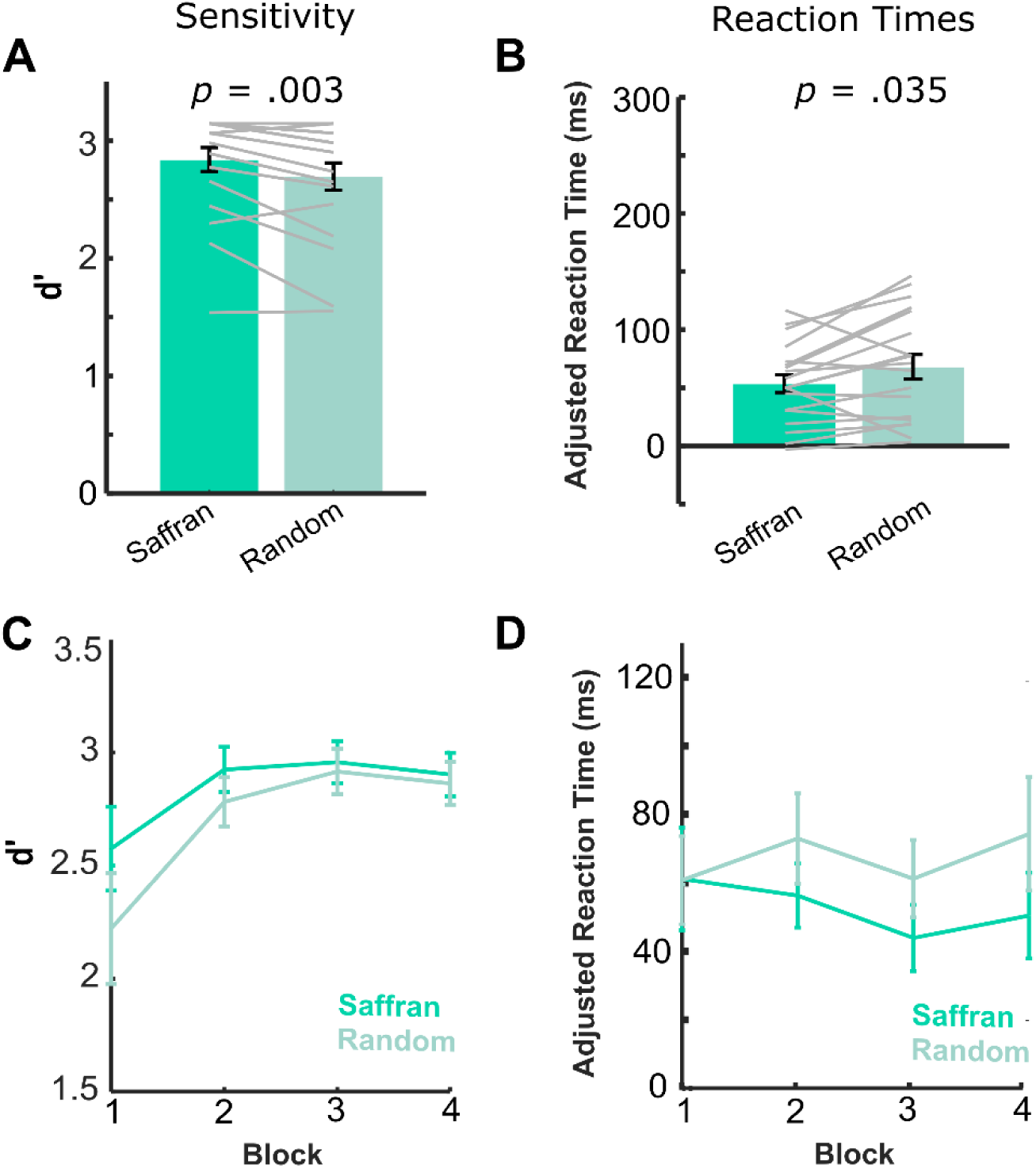
Results for Experiment 1 *Note.* The deviant detection task showed better performance for the Saffran sequences compared with Random sequences of tones. Sensitivity(d’) was significantly higher (**A**) and reaction times (**B**) were significantly shorter for Saffran relative to Random sequences. Reaction times were adjusted based on response time to a control condition. **C** and **D** show how responses changed across blocks. Gray lines indicate individual subjects. Error bars show ± SEM.

Reaction times did not significantly change with block (*F*(3.54) = .361, *p* = 0.706, *η_p_^2^* = .020), the effect of sequence type was present (*F*(1,18) = 5.224, *p* = .035, *η_p_^2^* = .225) with no interaction *F*(2,54) = .891, *p* = .403, *η_p_^2^* = .047, figure 2D).

These results suggest that performance on both measures was superior in the Saffran condition compared to the Random condition, indicating that the presence of a predictable structure enhances the detection of the deviant. Focusing on changes over time, for sensitivity to the deviant, the structural advantage is clearly evident from the first block. However, both sequence types show improved performance over time, and the likely ceiling effects by the fourth block suggest that the final block may not have been necessary. For reaction times, statistical effects did not differ over time, although graphically there is an indication that a reaction time advantage does not develop in the first block.

### Discussion

Experiment 1 investigated the influence of the probabilistic Saffran triplet structure on the detection of a deviant tone. Participants exhibited greater sensitivity and faster reaction times when detecting identical deviants within a Saffran sequence compared to a randomly ordered tone sequence, with the sensitivity effect emerging rapidly from the first block. These results indicate that, as has been observed for deterministic sequences, probabilistic sequences with a triplet structure facilitate the detection of a deviant. These findings support the idea that deviants become more saliant in more predictable contexts due to an increase in the prediction errors in high relative to lower precision/certainty sequences (Southwell & Chait, 2018).

However, the Saffran structure has limited complexity as it relies on adjacent probabilities; it also contains chunks of information that are deterministic in nature, since items within a triplet always co-occur. To explore whether these effects generalize to other statistical structures and vary systematically with uncertainty, experiment 2 was conducted.

## Experiment 2

Experiment 2 used the same deviant detection task, but the structured sequences were generated using a community structure (Benjamin et al., 2023; Karuza et al., 2017, 2019; Lynn et al., 2020; Schapiro et al., 2013). This structure, depicted in Figure 3 (top panel), contains 15 nodes; each node is connected to four other nodes and these interconnections are structured so as to group the nodes into three “communities”. If one considers a random walk through this structure, i.e., one that is only restricted by the available connections, there is both a high degree of uncertainty (each node has four possible options), but also a high chance of revisiting a node in the same community since these nodes are more interconnected. Based on previous research, it appears that the increased likelihood of revisiting a node within a community leads the perceiver to generate representations of the three communities. Importantly, this type of community structure cannot be learned based on transitional probability information alone, as the transitional probabilities among all stimuli are equal.

Participants were randomly assigned to one of three groups, with each group presented with stimuli that followed a particular traversal method based on Karuza et al. (2017): Unrestricted (termed “Random” in their article), Eulerian and Hamiltonian. Based on our analyses described below, these walks differed in terms of the resulting levels of uncertainty, with uncertainty levels decreasing from Random to Eulerian to Hamiltonian walks.

We hypothesized that participants would be sensitive to the underlying structure and show increased sensitivity and faster reaction times to the predictable sequences, as observed for Experiment 1. We also predicted that the results would be influenced by the differences in uncertainty, with the greatest advantage in the Hamiltonian condition and least advantage in the Unrestricted condition.

### Methods

#### Sample Size Selection

For this experiment, within group measures contrasting structure and random sequences could be based on experiment 1, and therefore 20 participants would be sufficient. However, our additional goal was to test the within-between interaction of the within-subject variable of sequence type (Structured vs Random) and between-subject variable walk (Unrestricted, Eulerian, Hamiltonian). A priori power analysis was computed for a repeated-measures, within-between interaction using G*Power software (Faul et al., 2007). Since we had no prior comparable test to estimate effect sizes, we used an effect size of, *η_p_^2^* = 0.02 (reasoning that such a small effect size should allow us to observe an effect if one is present) with α= .05; β = .8, giving a sample size of 42 participants per condition.

#### Participants

In total, 140 individuals were recruited, nine participants were excluded due to equipment issues, and one participant failed the control condition. The remaining participants were distributed as follows: 42 participants for the Random walk (Age: *M = 26.3, SD* = 6.6; males = 33, females = 9), 45 participants for the Eulerian walk (Age: *M = 26.0, SD* = 5.5; males = 8, females = 37), and 43 participants for the Hamiltonian walk (Age: *M = 25.4, SD* = 7.3; males = 18, females = 22). Initially, participants were assigned to a specific walk using the Gorilla balanced randomizer. After the exclusion of participants, individual experiment releases for each walk were required to replace the excluded subjects. Renumeration and ethics were the same as for Experiment 1.

#### Stimuli

Each trial contained one sequence of 96 tones lasting 4800ms. To avoid participants relying on low-level frequency cues to identify different communities, the 15 tones were arranged in descending order and divided into five groups, each comprising three frequencies. Within the graph structure, one tone from each group was randomly assigned to the three communities. As for experiment 1, five stimulus sets were used to further prevent the effect of low-level cues. The following sequence types were generated by traversing the community structure in different ways:

1. Unrestricted walk: A random starting node (tone) was selected from any community. Each tone could transition to any other tone connected to it within the graph structure. This process was repeated until the desired sequence length was achieved. Since there were no constraints, the minimum duration spent within a community could range from 0 to infinity. For instance, a walk might never reach a specific community, visit it for only one node, or, conversely, remain in a single community indefinitely. In our stimuli, the time spent within a community ranged from 1 to 75 nodes. The median duration was 10 nodes, with an average of 9 community transitions occurring during a 96-tone trial.
2. Eulerian walk: A random start node was chosen from any community, and all connections in the graph structure were traversed exactly once before returning to the start node. This process continued for multiple circuits of the structure until the sequence length was reached. The specific route taken for each circuit was random, with the constraint that a connection could not be repeated. This walk always results in a presentation of 30 tones per circuit. 10 nodes are visited in a community before moving to the next community (excluding the first community that is visited, as this is a random start point) with three transitions per circuit, on average the 96 tone trial will contain 9 community transitions.
3. Hamiltonian walk: A random start node was chosen from any community, and the structure was traversed in such a way that each node was encountered exactly once before returning to the start node. This process was repeated for the length of the sequence. Similar to the Eulerian walk, the exact route taken for each circuit was random, but with the restriction that a node could not be revisited within the same circuit. This walk always results in a presentation of 15 tones per circuit, visiting the five nodes in a community once, and with three community transitions per circuit. Each 96 tone trial will contain approximately 19 community transitions.

These three types of walks were selected because they were expected to differ in terms of uncertainty (Karuza et al., 2015). To quantify this uncertainty, a statistical learning model based on Prediction by Partial Matching (PPM) was used (Pearce et al., 2010). This model uses unsupervised statistical learning to estimate the uncertainty of an event prior to its occurrence by implementing a variable-order Markov model, where the amount of context used for prediction varies adaptively. The output of the model provides the information content (IC) for each tone, represented as the negative logarithm of the tone’s probability (-log P). Higher IC values indicate greater unexpectedness of the tone. The sequences in Experiment 1 have previously been modelled using this method (see Milne et al., 2021).

The IC values were separately computed for each of the three walks. Figure 3B displays the IC values for individual items within a sequence, averaged across sequences within a single stimulus set (without considering preceding trials). Figure 3C depicts the overall IC values across the entire experiment, computed as the average IC for each trial. The grey lines represent the IC values for the random sequences in both comparisons, showing that uncertainty remains high in random sequences since they cannot be predicted. In contrast, the uncertainty decreases for all predictable sequences, with the Hamiltonian Walk resulting in the greatest reduction in uncertainty and the Unrestricted walk resulting in the least reduction.

**Figure 3.**
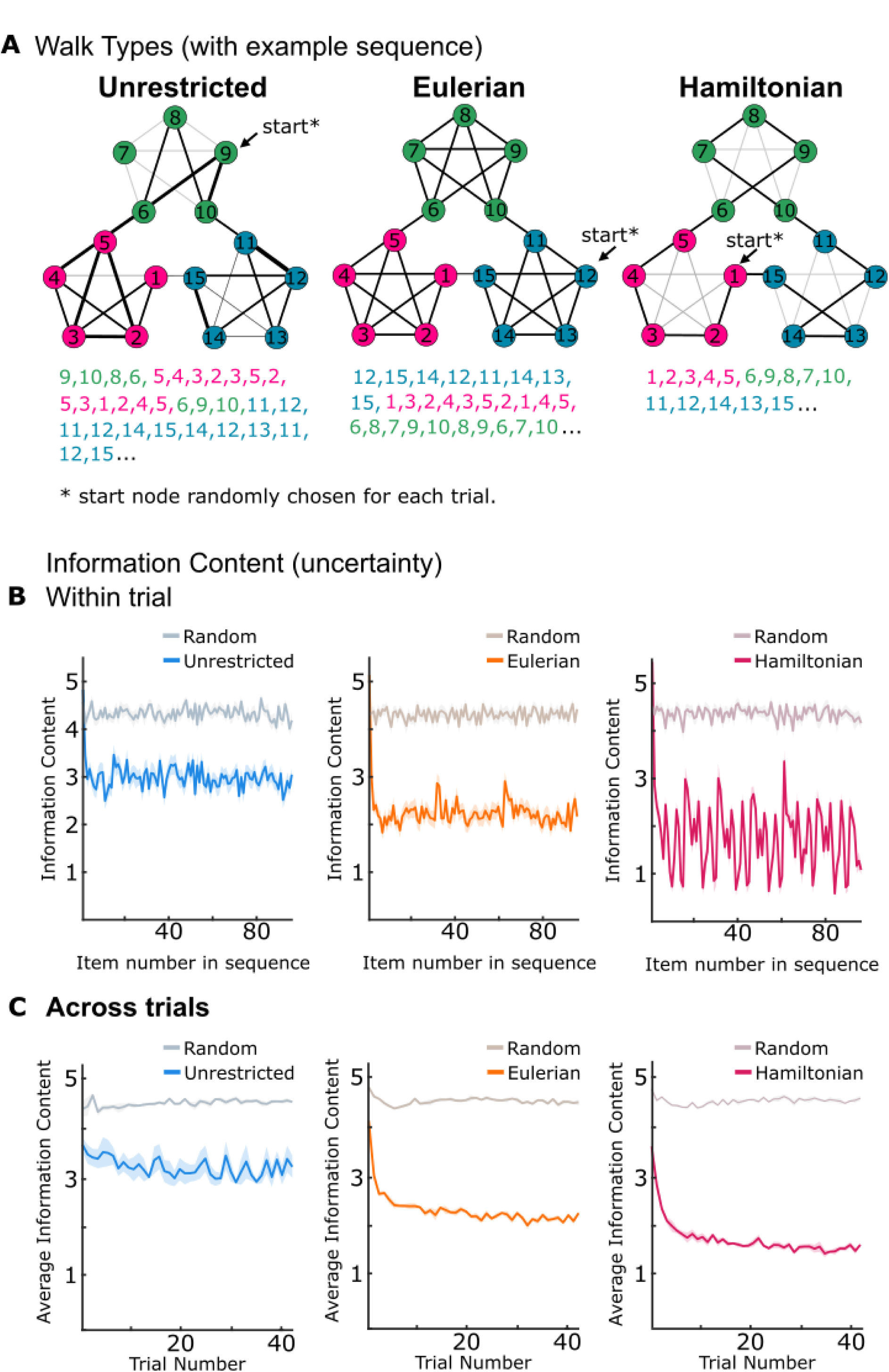
Stimuli design for Experiment 2 *Note.* Structured sequences were generated according to the graph structure in **A.** In this structure, each node connects to four other nodes; however, it is possible to separate the nodes into three communities (depicted by different colors), because nodes within a community are more interconnected. Each node is assigned a different tone frequency. Participants were separated into three conditions that varied based on how the structured sequence traversed the graph structure. Examples are shown in **A:** 1) Unrestricted walks (**left**) were generated by traversing the graph without any restrictions. The number sequence below the graph provides an example sequence using the node numbers and community colors. The black lines on the graph show connections that are used in the example; the thickness of the line increases as that connection is revisited. Gray lines are connections that are not visited in the example sequence; 2) Eulerian ( **middle**) walks visited every connection exactly once before beginning a new circuit of the graph; 3) Hamiltonian walks (**right**) visited every node exactly once before beginning a new circuit of the graph. These are based on the walks used by Karuza et al., (2017). The uncertainty for each walk was estimated using an ideal observer model (Pearce et al., 2015). Plotted in **B** and **C** is the information content (IC), the negative log probability (-log P); the higher the IC value the more unexpected the tone. **B** shows IC for a trial (averaged over 42 sequences for a single stimulus set). **C** shows IC across the experiment calculated by taking the average IC for each trial (averaged over the five stimulus sets). The IC to random sequences (gray hues) remains high for all walks. For structured conditions (colors), IC drops quickly, for individual trials and across the experiment. The Hamiltonian walk has the lowest average IC and the Unrestricted walk the highest. Without the accumulation of information from previous trials, there is more variability in IC for individual items in Hamiltonian sequences than the other two walks, likely due to each item being presented a single time per circuit. Shading indicates ± SEM.

#### Procedure

The deviant detection task followed the same procedure as for Experiment 1. Before beginning, participants were assigned to one of the three walks. All participants completed the same practice phase but the stimuli in the main phase differed based on the participants’ assigned walk. For all walks, there were three blocks, each consisting of 46 trials. Within each block, there were 21 structured sequences and 21 random sequences, 14 of which contained deviant tones. Additionally, there were four control trials. We opted to use only three blocks, a decision informed by evidence from Experiment 1, suggesting that the fourth block contributed little additional information. Our stimulus modelling also indicated that by the third block, there were only minor changes in information content for the different walks.

#### Data analysis

Sensitivity scores and reaction times were computed as described in Experiment 1. The main analysis of interest was a repeated-measures (RM) ANOVA with walk (Unrestricted, Eulerian, Hamiltonian) as a between-subjects factor and within-subject factors of sequence type (Structured, Random) and block (1,2,3). Where necessary, interactions involving walk were conducted by computing the structural advantage (Structure sequence – Random sequence) for each walk type. This was then subject to a one-way ANOVA and subsequent pairwise tests (Bonferroni corrected). These were normally distributed. Again, measures were computed for each block and averaged prior to statistical analysis. As there were 14“target” trials and 7 “non target” trials per condition the d’ ceiling was 3.27.

We also included planned comparisons to confirm the presence of the structure effect for each walk, to replicate the findings of Experiment 1. Due to non-normal data distribution Wilcoxon signed-rank tests were used.

### Results

#### Sensitivity (d’) to deviants

Participants were presented with sequences generated using an Unrestricted, Eulerian or Hamiltonian walk through the community structure. The detection of deviants was compared for sequences that followed one of these walks (Structured) relative to randomly ordered sequences of tones (Random). Planned comparisons confirmed the presence of the structure effect for each walk; Wilcoxon signed-rank tests showed significant differences for Unrestricted (*Z* = -2.950, *p* = .003, *d_z_* = 1.02), Eulerian *(Z* = -4.529, *p* < .001, *d_z_* = -1.85), and Hamiltonian (*Z* = -4.908, *p* < .001, *d_z_* = -2.25) walks (Figure 4A). Some participants reached the ceiling of 3.27.

**Figure 4.**
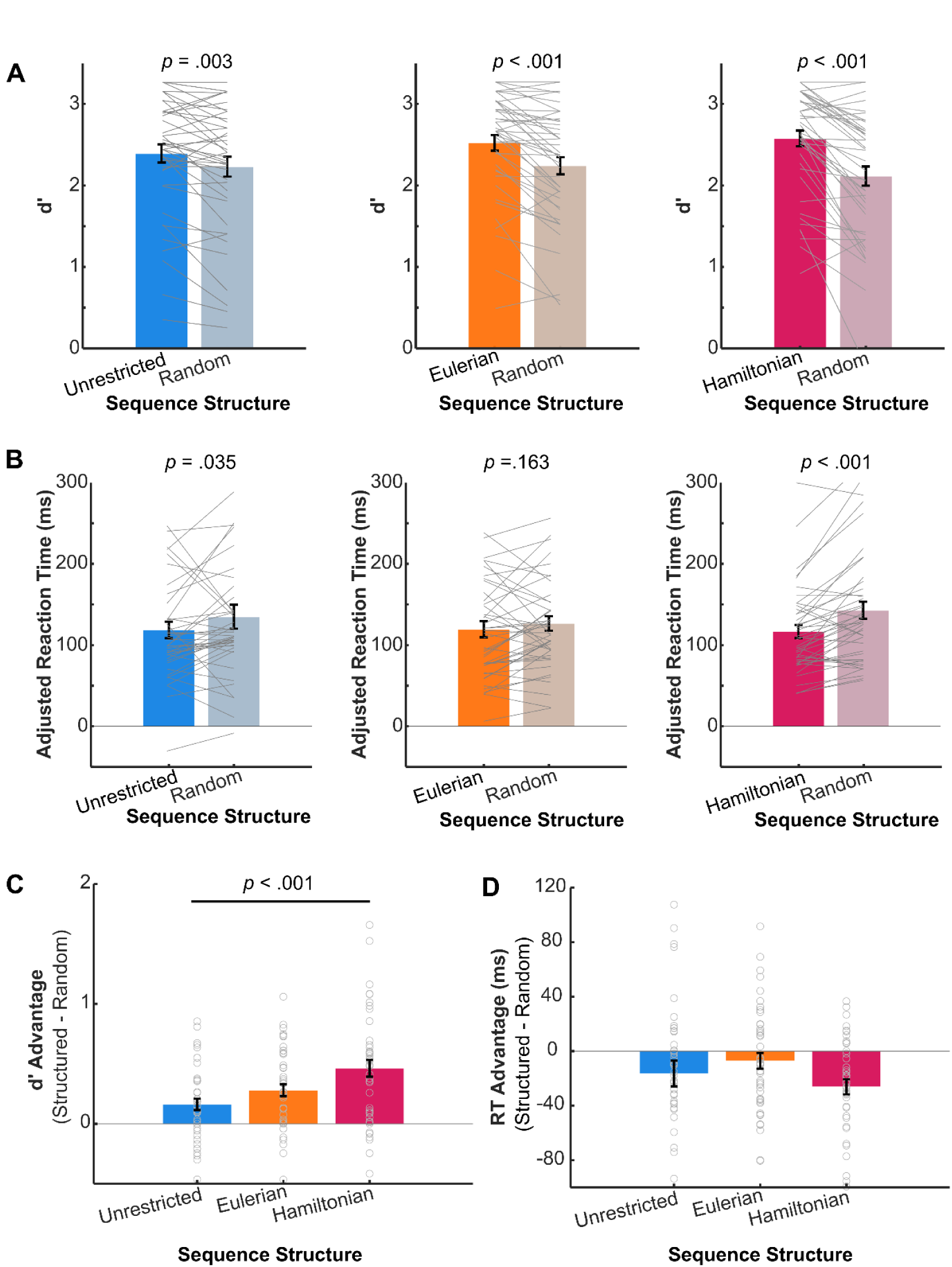
Main results for Community structure experiment (experiment 2). *Note.* There was greater sensitivity (d’) to the deviant in the structured compared to a random sequence of tones for all walks (**A, right-left:** Unrestricted, Eulerian, Hamiltonian) and faster reaction times for the Unrestricted and Hamiltonian walks (**B**). d’ results showed an interaction with walk type, with a greater impact of structure for the Hamiltonian than Random walk (**C**, **left**). Reaction times showed no interaction between sequence and walk (**C, right**). Grey lines/ circles represent individual participants. Error bars show ± SEM.

The RM ANOVA showed no significant main effect of walk (*F*(2,127 = .116, p = .891, η_p_^2^ = .0232). The lack of a main effect of walk is important as it suggests comparable levels of general performance across the different participant groups. There was, however, a significant main effect of sequence type (Structured vs Random: *F*(1,127) = 85.614, *p* <.001, *η_p_^2^* = .403) and a main effect of block (*F*(2,254) = 41.503, p <.001, η_p_^2^ = .397). Regarding the main effect of block, pairwise post hoc tests revealed that performance on each subsequent block was significantly better than the previous block (block 1 vs 2: *p* < .001, CI [-.39 -.14]; block 2 vs 3: p = .001, CI [-0.27 -0.05]).

There was a significant interaction of sequence type with walk (*F*(2,127) = 7.088, *p* = .001, *η_p_^2^* = .100; figure 4A) but no significant interactions between block and walk or block and sequence type, and no significant three-way interaction (p-values > .1, for block data see figure 5A). Investigation of the interaction of sequence type and walk using the structural advantage for each walk type (Figure 4C, left) found a significant effect of walk (*F*(2,127) = 7.088, *p* = .001) driven by significantly larger advantage for the Hamiltonian walk compared to Unrestricted walk (M difference = 0.30, SEM = 0.08, *p* = .001, 95% CI [0.11, 0.50]). The structural advantage for Eulerian sequences did not significantly differ from Hamiltonian (M difference = -0.18, SEM = 0.08, *p* = .069, 95% CI [-0.38, 0.01]) or Unrestricted sequences (M difference = 0.12, SEM = 0.8, *p* = .420, 95% CI [-0.08, 0.31], figure 4C).

**Figure 5.**
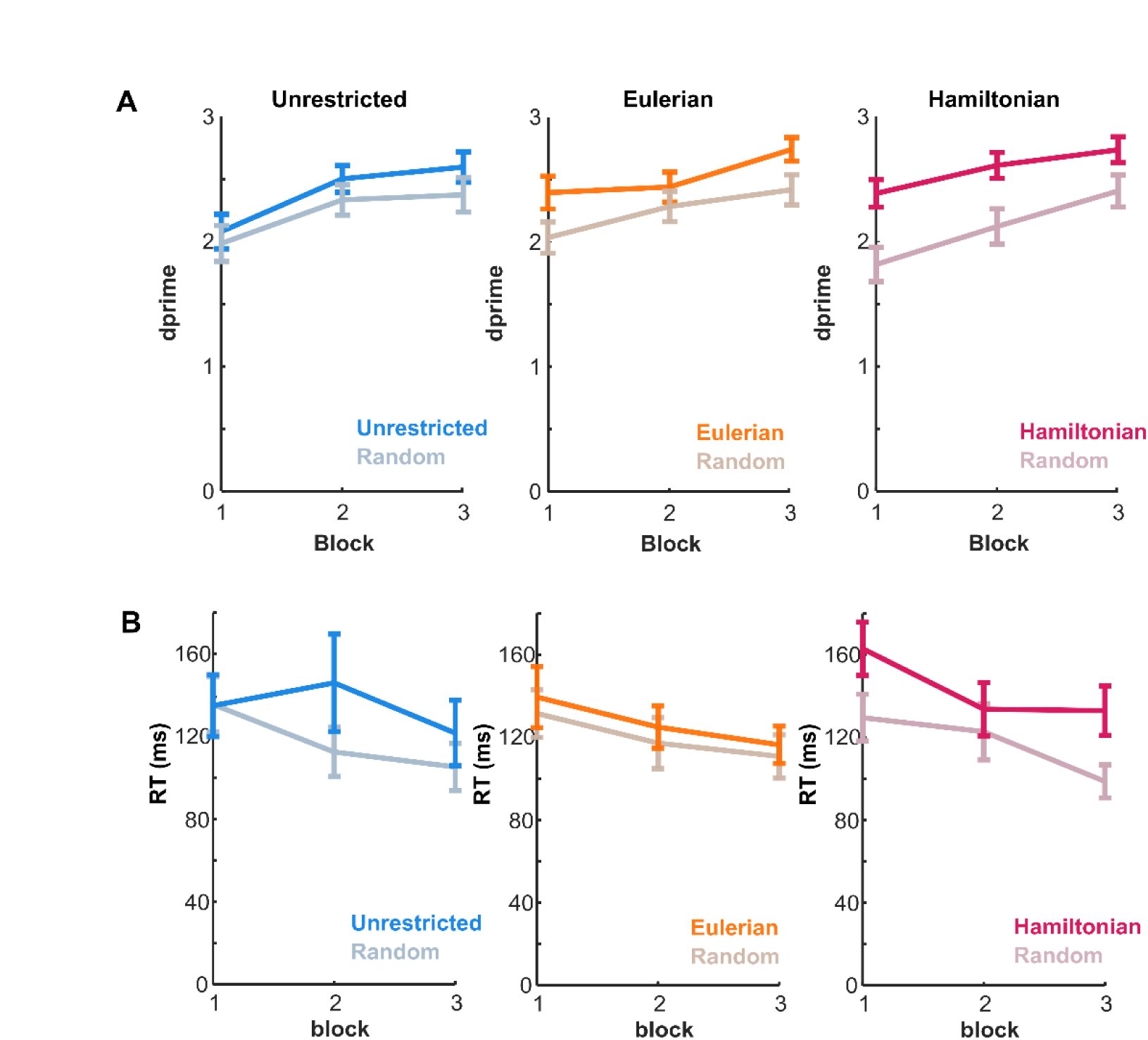
Experiment 2 results across blocks *Note.* Sensitivity (**A**) and reaction times (**B**) improved over time but there was no interaction with sequence or walk. Error bars show ± SEM.

### Reaction times

Planned comparisons showed that participants were significantly faster on structured sequences for Unrestricted (*Z* = -2.107, *p* = .035, *d_z_* = -0.70) and Hamiltonian (*Z* = -3.912, *p* < .001, d_z_ = -1.5) walks, but not for Eulerian walks (*Z* = -1.394, *p* = .163, *d_z_* = -0.43).

The main effect of walk was not significant (*F*(2,127 = .116, p = .891, η_p_^2^ = .0232), again suggesting overall comparable performance across groups. We observed a significant main effect of sequence type *F*(1,127) = 16.469, p <.001, η_p_^2^ = .115, figure 4B). There was also a significant main effect of block (*F*(2,254) = 41.503, p <.001, η_p_^2^ = .397), although this time post hoc tests showed that the effect was driven by a significant difference between blocks 1 and 3 (*p* <.001, 95% CI [10.45 38.94] ) but not 1 and 2 (*p* = .117, 95% CI [-2.10 27.72]) or 2 and 3 (*p* = .068, 95% CI [-0.60 24.34]), suggesting a more subtle improvement over time. There were no significant interactions with Block (*F*(2,254) = .126, p = .874, η_p_^2^ = .001, nor a three way interaction between block, sequence and walk (*F*(2,254) = 1.259, *p* = .287 , η_p_^2^ = .019; figure 5B).

In summary, the results from Experiment 2 were consistent with those of Experiment 1, demonstrating that structured sequences led to better deviant tone detection and faster reaction times compared to random sequences. The walk used to generate the structured sequences impacted performance, with significantly better sensitivity to deviants in the Hamiltonian than Unrestricted condition. The block analysis indicated significant improvements in performance and reaction times over time, suggesting a practice effect, but this did not interact with the sequence or walk types.

### Discussion

Experiment 2 aimed to determine if the facilitation observed in Experiment 1, where the presence of the Saffran structure improved deviant detection, would generalize to a more complex statistical structure. Using sequences with an underlying community structure, we found robust facilitation of deviant detection, characterized by greater sensitivity and faster reaction times in structured compared to random sequences. Experiment 2 also manipulated how the community structure was traversed to create varying levels of uncertainty. Consistent with our hypothesis that sensitivity to deviants would scale with uncertainty, deviant detection was best for the walk associated with the least uncertainty; however, there were no differences in reaction times. These effects showed no interaction with time, as performance improved over blocks for both structured and random sequences but remained consistent across walks. Note that we only sought sufficient power for the main interaction between sequence type and walk, the focus of this experiment, and our sample may not have been large enough for the additional within-subject interactions; therefore we do not draw conclusions from the absence of any interaction with block.

Critically, even the unrestricted walk, which represents an exceedingly complex statistical structure, was associated with a “predictability advantage”, revealing listeners attunement to statistical regularities even in rapid sound sequences where deliberate tracking of structure is not possible.

In both this experiment and the previous one (Experiment 1), the stimulus sequences used may have elicited some degree of auditory streaming—a cognitive process where the auditory system organizes incoming sounds into perceptual groups or “streams” based on features like frequency (Moore & Gockel, 2012). Theoretically, streaming could interfere with listeners’ ability to detect adjacent relationships between items by segregating sounds into separate streams.

However, several aspects of the present stimuli suggest that auditory streaming did not significantly impact the observed findings. First, the same frequency pool used here was employed in previous studies (Barascud et al., 2016; Bianco et al., 2020; Harrison et al., 2021; Hu et al., 2024; Southwell & Chait, 2018; Zhao et al., 2025), which demonstrated robust effects consistent with listeners learning sequential relationships, even in fast sequences (e.g., 20 Hz) similar to those used here. Using modelling, Zhao et al. (2024) demonstrated that behavioural performance aligned more closely with a model that tracks the entire sequence using a single Gaussian prior, rather than a listening strategy that divides the sequence into channels—a process akin to perceptual streaming.

Most importantly, the same frequency structure was applied to both the random and structured sequences in the present study. This ensures that both sequence types were equally susceptible to any potential streaming effects. Consequently, any observed advantage of structured sequences over random ones is indicative of listeners having successfully acquired sequential statistics, rather than being confounded by streaming-related artifacts.

Furthermore, the community structures were designed so that the tones allocated to each community always included a random. mix of high and low frequencies, with their positions within the community randomly assigned. This design minimizes the likelihood of specific frequency-related segregation impacting the results.

Finally, we used multiple stimulus sets with varied tone allocations, reducing the possibility of consistent non-adjacent, acoustically-based statistical relationships influencing the outcomes. These measures collectively bolster confidence that the findings reflect true differences in listeners’ ability to learn and utilize sequential statistics, rather than artifacts introduced by auditory streaming.

## Experiment 3

Experiment 2 confirmed the presence of a structural effect in sequences generated using the community structure and found that variations in uncertainty between different walks influenced the ability to accurately and quickly detect deviant tones. Experiment 3 aimed to further explore these effects, specifically testing if fine-grained structural information would modulate deviant detection.

In serial response time studies using visual versions of the community structure paradigm, a delay in response time at community boundaries has been observed for Eulerian and Random walks (Karuza et al., 2017). This “surprisal” response indicates that participants were more likely to expect an item from the same community than from a different community.

In Experiment 3, we hypothesized that the processes of shifting from the context of one community to another creates a perceptual boundary that will be associated with a momentary increase in uncertainty. As a result, listeners may rely less on their predictions than when within an established community and be less sensitive to deviants that occur within this period of uncertainty. To test this, we embedded deviants either right after a change in community or after the sequence had been within a community for at least five tones. We predicted better detection and faster reaction times to deviants if they appeared within a community. We used only the Eulerian walk, as based on Karuza et al., (2017), there should be a robust effect of community structure, and the more uniform nature of this walk allowed for easier placement of the deviants in the two conditions.

### Method

#### Sample Size Selection

Using the sample size from Experiment 2 and α= 0.05; β = 0.8 in G*Power software (Faul et al., 2007) with a sensitivity power analysis for a repeated-measures ANOVA, with one group and three measurements (correlation of 0.5 and non-sphericity correction of 1), indicated that we would observe a significant effect at an effect size of η_p_^2^ = 0.036. We were satisfied that effects smaller than this were unlikely to be of relevance and therefore this sample size should be sufficient to detect an effect if one is present. We sought an extra 15% of cases, a total of 52 subjects, to provide sufficient power for the use of non-parametric tests (Lehmann, 1998).

#### Participants

Fifty-seven participants were recruited; five participants were excluded prior to analysis due to using a non-windows operating system, leaving a final sample of 52 (Mean age = 25.8, SD = 5.2 years; 11 female, 41 male).

#### Stimuli

The Eulerian, Random, and Control sequences were generated in the same manner as in Experiment 2. Additional constraints were applied when introducing the deviants to create conditions where the deviant was either at the boundary of a community or within a community. For “Boundary” deviants, we identified points where the sequence exited one community and entered a new one. The deviant was then placed immediately after the first node of the new community, node labelled B in figure 6. For “Within” community deviants, there had to be at least four transitions within a community before the deviant could occur; the W on figure 6 indicates the earliest location within community deviants could occur relative to a community boundary. To ensure sufficient time to establish the structure, the deviant could not occur until after one full circuit of the structure, which was 30 tones/1500ms. The deviant could also not occur in the final 500ms. The remaining time window was divided into three equal windows, and the deviants were evenly distributed across these for each condition. Four stimulus sets were generated to minimize the impact of low-level auditory cues.

**Figure 6.**
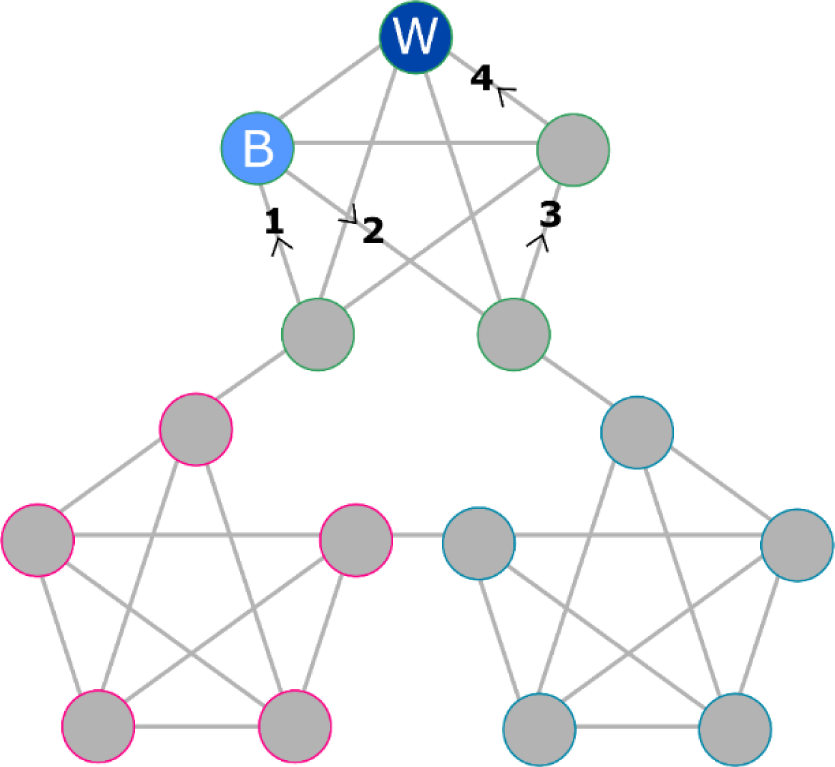
Experiment 3 deviant positioning *Note.* Deviants were placed either following a community boundary or within an established community. A boundary deviant always occurred exactly one tone after entering a new community, example marked with a B (“Boundary”). Within community deviants could only occur after at least four within community transitions, for example, in the position marked with W (“Within). Only sequences generated from the Eulerian walk were used in this experiment and only one type of deviant could occur per trial. Random and control conditions were also included.

#### Procedure

Experiment 3 followed the same procedure as Experiment 2 but included only the Eulerian, Random, and Control walks, not the Hamiltonian or Unrestricted. As in the previous experiment, there was a practice phase followed by three blocks. Each block consisted of 52 trials with the following conditions: 12 Eulerian sequences containing a boundary deviant, 12 Eulerian sequences containing a within community deviant, 12 Random sequences containing a deviant, 6 Eulerian sequences without a deviant, 6 Random sequences without a deviant, and 4 control trials.

#### Data Analysis

Reaction times were computed as described in Experiments 1 and 2. It was not appropriate to calculate performance scores using dprime as the two deviant conditions would be measured against the same non-deviant condition. Therefore, we analyzed hit rate for the three deviant conditions and separately included false alarm analysis for the Eulerian vs Random non-deviant trials. All measures violated normality and therefore non-parametric Friedman tests were used to compare performance across the three deviant conditions (Eul-Boundary, Eul-Within, Random). Pairwise/post hoc analyses used Wilcoxon signed-rank tests with Bonferroni corrections applied.

## Results and Discussion

There was a significant effect of deviant condition on hit rate (*ꓫ^2^* (2)= 14.11, *p* = .001, *W*= .136), which was driven by a significant difference between the Eulerian and Random conditions (Eul-boundary vs Random: *Z* = -3.197, *p*= .003, *d_z_* = -0.98; Eul-within vs Random: *Z* = -4.05, *p* < .001, *d_z_* = -1.35; Figure 7A) but there was no significant difference when comparing the two Eulerian conditions (*Z* = -.150, *p* = 0.402, *d_z_* = -0.42). The same pattern was observed for the reaction times (Figure 7B): a significant main effect of condition (*ꓫ^2^* (2) = 14.106, *p* <.001, *W* = .193), with significant faster reaction times for deviants in the Random sequences relative to the deviants in the Eulerian sequences, regardless of whether they were a community boundary (*Z* = -4.034, p <. 001, *d_z_ =* -1.35) or within a community (*Z* = -2.124, *p* = .018, *d_z_* = -0.61). There was no difference based on deviant position in the Eulerian sequences (*Z* = -1.202, *p =* .687, *d_z_ =* -0.35). The false alarm rate compute from non-deviant trials was significantly higher in the Random compared to Eulerian walk (*Z* = - 5.124, p < .001, Figure 7C, *d_z_ =* -2.02) mirroring the d’ results of Experiment 2.

**Figure 7.**
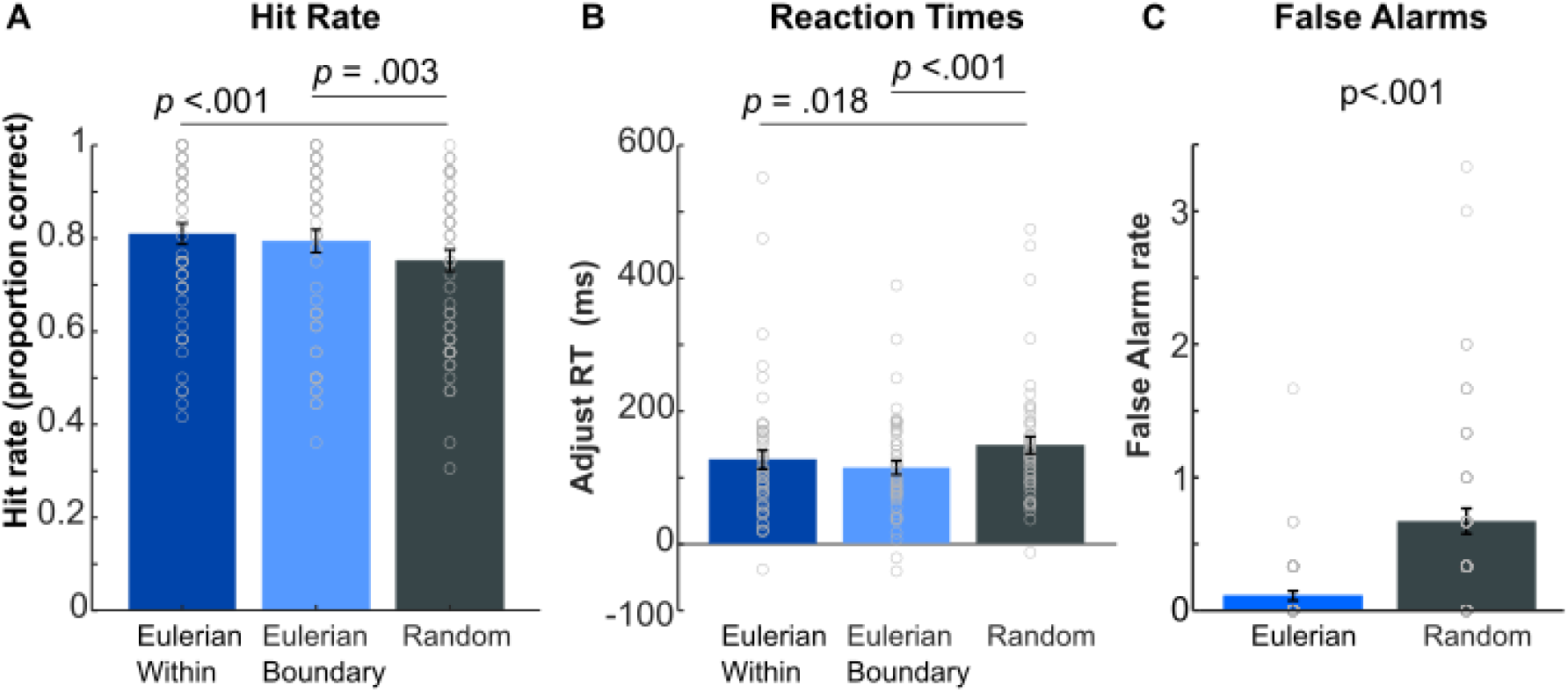
Experiment 3 results Sensitivity (**A**) and reaction times (**B**) to deviants in Random sequences and Eulerian sequences where the deviant was positioned either within a community or immediately after the boundary of two communities. (**C**) False alarm rate for non-deviant trials. There were only significant differences between Eulerian and Random sequences. Grey circles represent individual participants. Error bars show ± SEM.

In sum, these findings further support the idea that general predictability derived from the underlying community structure enhances the detection of deviant tones. However, our results found that proximity to community boundary did not significantly impact deviant detection. Since this is a null result, it does not provide evidence that there is no difference between these conditions but in the next section we discuss why no significant difference may have been observed.

## General Discussion

Over three experiments, we investigated the influence of sequence predictability on the detection of an unexpected deviant tone. The results demonstrated that sequences that followed a probabilistic structure enhanced the detection of a tone of an outlying frequency, relative to the same tones presented in a random order. Behavior was even facilitated in the “Unrestricted walk” variant of the community structure stimuli (Experiment 2) revealing listeners’ readiness to discover complex predictable structure (each node connected to 4 other nodes) in rapidly unfolding sound sequences. Furthermore, deviants were more salient in sequences that could be more accurately predicted, i.e., had lower uncertainty. Therefore, this suggests that violation signals used to detect the deviant may be sensitive to the reliability of the available statistical information. These findings open the door to further understand the mechanisms underlying efficient processing of sensory input.

The data presented here are based on key press responses collected remotely via online platforms, yet the findings mirror those from neural (EEG) recordings, specifically that the variability of “standards” in an oddball paradigm reduces responses to identical deviant tones (Daikhin & Ahissar, 2012; Winkler et al., 1990; Furl et al., 2011; Garrido et al., 2013; Southwell & Chait, 2018). At a methodological level, these results provide evidence that the extraction of underlying probabilistic cues can be behaviorally probed using simple detection violation tasks. Therefore, this rapid, simple experiment has the potential to serve as a behavioral lens into the brain’s ability to extract predictability from various statistical structures and the mechanisms used during sensory processing. We review the implications of the main findings below.

### Mechanisms for predictability-modulated deviant detection

We hypothesized that the deviants would be better detected in structured relative to random sequences of tones, based on the theory that listeners are continuously generating an internal predictive model of sensory input. The accuracy of this model is continuously monitored via prediction errors that indicate the discrepancy between the model and sensory input. In structured sequences, where that context can be relied on, listeners upweight prediction errors and are highly sensitive to changes in their value; as a result, they should be able to rapidly detect unexpected events (Feldman & Friston, 2010; Friston & Kiebel, 2009; Heilbron & Chait, 2018; Press & Yon, 2019; Yon & Frith, 2021).

Our findings should also be considered alongside other mechanisms, though not mutually exclusive with predictive coding, that influence how we process and prioritize sensory input. For example, regularity has been found to increase the likelihood that a series of tones are represented as a unified perceptual object rather than just a stream of individual items, i.e. perceptual binding (Andreou et al., 2011; Winkler et al., 2009). In this scenario the appearance of a deviant tone may be enhanced not only by the upweighting of prediction errors, but also because it is treated as a separate perceptual object. Perceptual binding could also affect the resources required to process the sensory stream. If, unlike the random sequences, structured sequences are treated as an identified stable source of information, the listener may more easily suppress or ignore the content (De Lange et al., 2018; Itti & Baldi, 2009), thus freeing cognitive resources to detect unexpected changes in the environment (a similar point is also made by Planton et al., 2023). This is consistent with evidence that when presented concurrently (dichotically), unpredictable rather than predictable sequences capture attention (Southwell & Chait, 2018). For the present stimuli, it is also possible that if listeners are actively monitoring for the deviants while attempting to suppress the main sequence, e.g. supressing central frequencies to attend to the extreme edges, this process may be easier for structured sequences compared to random ones.

A similar effect would also be observed if predictability is facilitating sensory processing of the main sequence, i.e. impacting cognitive effort. Milne et al., (2021) demonstrated that as the listener establishes the presence of both deterministic patterns and a probabilistic Saffran structure, there is a decrease in pupil size relative to unstructured sequences. Pupil size is modulated by the neural systems that regulate effort and attention, with larger pupil sizes associated with greater effort (Ohlenforst et al., 2018; Zekveld et al., 2018). Thus, the reduction in pupil size is consistent with evidence that predictability can facilitate processing and conserve cognitive effort. Similarly, in the predictive coding framework, driving the shift in cognitive effort may be a switch from resource-demanding bottom-up processing required to process random sequences to more efficient top-down processing when the listener can instead match incoming tones to a predictive model (Friston & Kiebel, 2009).

Critically, all these explanations for the observed differences between the structured and random sequences hinge on listeners having successfully acquired the predictable structure inherent in the stimuli. They diverge, however, in how they propose this acquired knowledge is utilized during the process of deviance detection.

Some accounts may suggest that the predictable structure enhances anticipation, allowing listeners to more efficiently identify deviations from the expected pattern. Other perspectives might emphasize a reduced cognitive load in processing structured sequences, freeing resources for detecting irregularities. Alternatively, the structured sequences could facilitate suppression of irrelevant information, making deviant tones more salient in comparison. Each explanation reflects a distinct mechanism by which predictability interacts with auditory processing to influence deviance detection.

### The influence of predictability versus community structure

Experiment 3 aimed to investigate whether the response to deviant stimuli would be influenced by recent transitions between communities. Specifically, we compared responses to deviants occurring immediately after a transition to those occurring within a community. Previous studies (Benjamin et al., 2023; Karuza et al., 2017; Schapiro et al., 2013) have demonstrated that participants can consciously parse sequences into these communities, despite uniform transitional probabilities at the boundaries. Additionally, in a serial reaction time task, participants exhibited a response delay when encountering an item representing a transition to a new community (Karuza et al., 2017).

We hypothesized that a transition between communities, and presumably the process of shifting from the context of one community to another, constitutes a perceptual boundary associated with a momentary increase in uncertainty. Therefore, if participants in our experiment are representing the sequences as community structures, we would expect reduced performance at community boundaries. However, contrary to expectations, we observed no effect based on the position of the deviant relative to the community boundary. This may be interpreted to suggest that listeners did not parse the sequences based on the community structure but instead tracked the overall uncertainty associated with the sequence’s statistical structure. In the community structure, transitional probabilities are uniform across community boundaries, and it is possible that the violation signal may only track this characteristic.

Alternatively, it may be that listeners do represent community structure but that the higher-order structure does not influence the saliency of deviant stimuli, or that higher-order structures do in fact impact the processing of deviant stimuli, but the behavioral response is not sensitive enough to detect it. Further research, such as studying the neural response via EEG could help to disentangle these explanations and additionally elucidate the underling mechanisms and refine our understanding of how participants perceive and respond to complex sequence structures.

### General insights into statistical learning

Recent perspectives on statistical learning recommends that research should focus on statistical learning mechanisms that reflect real-world environments (Bogaerts et al., 2022), using methods appropriate for a given sensory modality and domain, rather than “generic tasks” (Bogaerts et al., 2022, p34). The current paradigm is based on methodologies, e.g. stimuli and techniques, that seek to understand fundamental components of auditory perceptual processing (Denham & Winkler, 2022; Heilbron & Chait, 2018; Khouri & Nelken, 2015). Furthermore, there are parallels with our day-to-day use of predictive information which is implicitly accumulated, and we would argue that a key function of extracting these cues is to ensure efficient navigation of the auditory environment and sensitivity to unexpected events. Therefore, the current data and paradigm provide crucial insights into auditory-focused statistical learning that can be drawn upon as the field seeks a unified theory of statistical learning that accounts for modality differences and domain-general phenomena.

## Conclusions

In the current study, we investigated humans’ sensitivity to predictability in rapid auditory sequences. Specifically, we used tones that were presented too rapidly for conscious tracking of the sequence structure. We found that predictability reliably modulates deviant detection, enhancing sensitivity and reaction times across two different structures and varying with the uncertainty of the predictable sequence in line with predictive coding frameworks. This study also marks the first attempt to employ a community structure with such rapid auditory stimuli. Beyond its immediate implications, this work lays the foundation for testing alternative manipulations of predictability via online platforms. Its simplicity and efficiency offer additional potential for probing individual differences, exploring deficits in how certain populations process rapid auditory information, and laying the groundwork for further investigation into the neural basis of how predictability modulates the brain’s response to unexpected events.

## Co-Author contributions

MC and CMC contributed to conceptualization, methodology and writing-reviewing and editing.

## Acknowledgements

This work was funded by a Wellcome Trust grant [213686/Z/18/Z] to AEM

## References

1. Al Roumi, F., Planton, S., Wang, L., & Dehaene, S. (2023). Brain-imaging evidence for compression of binary sound sequences in human memory. ELife, 12, e84376. 10.7554/eLife.84376

2. Andreou, L. V., Kashino, M., & Chait, M. (2011). The role of temporal regularity in auditory segregation. Hearing Research, 280(1–2), 228–235. 10.1016/j.heares.2011.06.001

3. Anwyl-Irvine, A. L., Massonnié, J., Flitton, A., Kirkham, N., & Evershed, J. K. (2020). Gorilla in our midst: An online behavioral experiment builder. Behavior Research Methods, 52, 388–407. 10.3758/s13428-019-01237-x

4. Appelbaum, M., Cooper, H., Kline, R. B., Mayo-Wilson, E., Nezu, A. M., & Rao, S. M. (2018). Journal Article Reporting Standards for Quantitative Research in Psychology. American Psychologist, 73(7).

5. Aslin, R. N., & Newport, E. L. (2012). Statistical learning from acquiring specific items to forming general rules. Current Directions in Psychological Science, 21(3), 170–176. 10.1177/0963721412436806

6. Barascud, N., Pearce, M. T., Griffiths, T. D., Friston, K. J., & Chait, M. (2016). Brain responses in humans reveal ideal observer-like sensitivity to complex acoustic patterns. Proceedings of the National Academy of Sciences of the United States of America, 113(5), E616–E625. 10.1073/pnas.1508523113

7. Batterink, L. J., & Paller, K. A. (2017). Online neural monitoring of statistical learning. Cortex, 90, 31–45. 10.1016/j.cortex.2017.02.004

8. Benjamin, L., Fló, A., Al Roumi, F., & Dehaene-Lambertz, G. (2023). Humans parsimoniously represent auditory sequences by pruning and completing the underlying network structure. ELife, 12, e86430. 10.7554/eLife.86430

9. Benjamin, L., Sablé-Meyer, M., Fló, A., Dehaene-Lambertz, G., & Roumi, F. Al. (2024). Long-Horizon Associative Learning Explains Human Sensitivity to Statistical and Network Structures in Auditory Sequences. Journal of Neuroscience, 44(14), e1369232024. 10.1523/JNEUROSCI.1369-23.2024

10. Bianco, R., Harrison, P. M., Hu, M., Bolger, C., Picken, S., Pearce, M. T., & Chait, M. (2020). Long-term implicit memory for sequential auditory patterns in humans. ELife, 9. 10.7554/eLife.56073

11. Bogaerts, L., Siegelman, N., Christiansen, M. H., & Frost, R. (2022). Is there such a thing as a ‘good statistical learner’? In Trends in Cognitive Sciences (Vol. 26, Issue 1). 10.1016/j.tics.2021.10.012

12. Conway, C. M. (2020). How does the brain learn environmental structure? Ten core principles for understanding the neurocognitive mechanisms of statistical learning. Neuroscience and Biobehavioral Reviews, 112, 279–299. 10.1016/j.neubiorev.2020.01.032

13. Daikhin, L., & Ahissar, M. (2015). Fast Learning of Simple Perceptual Discriminations Reduces Brain Activation in Working Memory and in High-level Auditory Regions. Journal of Cognitive Neuroscience, 27(7), 1308–1321. 10.1162/jocn_a_00786

14. Daltrozzo, J., & Conway, C. M. (2014). Neurocognitive mechanisms of statistical-sequential learning: what do event-related potentials tell us? Frontiers in Human Neuroscience, 8(437), 1–22.

15. De Lange, F. P., Heilbron, M., & Kok, P. (2018). How Do Expectations Shape Perception? Trends in Cognitive Sciences, 22(9), 764–779. 10.1016/j.tics.2018.06.002

16. Dehaene, S., Meyniel, F., Wacongne, C., Wang, L., & Pallier, C. (2015). The Neural Representation of Sequences: From Transition Probabilities to Algebraic Patterns and Linguistic Trees. Neuron, 88(1), 2–19. 10.1016/j.neuron.2015.09.019

17. Denham, S., & Winkler, I. (2022). Auditory Perceptual Organization. In D. Jaeger & R. Jung (Eds.), Encyclopedia of Computational Neuroscience (pp. 277–288). Springer New York. 10.1007/978-1-0716-1006-0_100

18. Deocampo, J. A., King, T. Z., & Conway, C. M. (2019). Concurrent Learning of Adjacent and Nonadjacent Dependencies in Visuo-Spatial and Visuo-Verbal Sequences. Frontiers in Psychology, 10, 436813. 10.3389/fpsyg.2019.01107

19. Ekman, M., Kok, P., & De Lange, F. P. (2017). Time-compressed preplay of anticipated events in human primary visual cortex. Nature Communications, 8, 15276. 10.1038/ncomms15276

20. Erickson, L. C., & Thiessen, E. D. (2015). Statistical learning of language: Theory, validity, and predictions of a statistical learning account of language acquisition. Developmental Review, 37, 66–108. 10.1016/j.dr.2015.05.002

21. Faul, F., Erdfelder, E., Lang, A.-G., & Buchner, A. (2007). G*Power 3: A flexible statistical power analysis program for the social, behavioral, and biomedical sciences. Behavior Research Methods, 39(2), 175–191. 10.3758/BF03193146

22. Feldman, H., & Friston, K. J. (2010). Attention, Uncertainty, and Free-Energy. Frontiers in Human Neuroscience, 4, 215. 10.3389/fnhum.2010.00215

23. Friston, K. (2005). A theory of cortical responses. Philosophical Transactions of the Royal Society of London B: Biological Sciences, 360(1456), 815–836.

24. Friston, K., & Kiebel, S. (2009). Predictive coding under the free-energy principle. Philosophical Transactions of the Royal Society B: Biological Sciences, 364(1521), 1211–1221. 10.1098/rstb.2008.0300

25. Frost, R., Armstrong, B. C., Siegelman, N., & Christiansen, M. H. (2015). Domain generality versus modality specificity: the paradox of statistical learning. Trends in Cognitive Sciences, 19(3), 117–125. 10.1016/j.tics.2014.12.010

26. Furl, N., Kumar, S., Alter, K., Durrant, S., Shawe-Taylor, J., & Griffiths, T. D. (2011). Neural prediction of higher-order auditory sequence statistics. NeuroImage, 54(3), 2267–2277. 10.1016/j.neuroimage.2010.10.038

27. Garrido, M. I., Sahani, M., & Dolan, R. J. (2013). Outlier Responses Reflect Sensitivity to Statistical Structure in the Human Brain. PLoS Computational Biology, 9(3), e1002999. 10.1371/journal.pcbi.1002999

28. Girvan, M., & Newman, M. E. J. (2002). Community structure in social and biological networks. Proceedings of the National Academy of Sciences of the United States of America, 99(12), 7821– 7826. 10.1073/pnas.122653799

29. Harrison, A. W., Mannion, D. J., Jack, B. N., Griffiths, O., Hughes, G., & Whitford, T. J. (2021). Sensory attenuation is modulated by the contrasting effects of predictability and control. NeuroImage, 237, 118103. 10.1016/J.NEUROIMAGE.2021.118103

30. Heilbron, M., & Chait, M. (2018). Great Expectations: Is there Evidence for Predictive Coding in Auditory Cortex? Neuroscience, 389, 54–73. 10.1016/j.neuroscience.2017.07.061

31. Herrmann, B., Henry, M. J., Fromboluti, E. K., McAuley, J. D., & Obleser, J. (2015). Statistical context shapes stimulus-specific adaptation in human auditory cortex. Journal of Neurophysiology, 113(7), 2582–2591. 10.1152/jn.00634.2014

32. Hu, M., Bianco, R., Hidalgo, A. R., & Chait, M. (2024). Concurrent Encoding of Sequence Predictability and Event-Evoked Prediction Error in Unfolding Auditory Patterns. Journal of Neuroscience, 44(14), e1894232024. 10.1523/JNEUROSCI.1894-23.2024

33. Itti, L., & Baldi, P. (2009). Bayesian surprise attracts human attention. Vision Research, 49(10), 1295– 1306. 10.1016/j.visres.2008.09.007

34. Kanai, R., Komura, Y., Shipp, S., & Friston, K. (2015). Cerebral hierarchies: predictive processing, precision and the pulvinar. Philosophical Transactions of the Royal Society B: Biological Sciences, 370(1668), 20140169. 10.1098/rstb.2014.0169

35. Karuza, E. A., Kahn, A. E., & Bassett, D. S. (2019). Human sensitivity to community structure is robust to topological variation. Complexity, 2019, 8379321. 10.1155/2019/8379321

36. Karuza, E. A., Kahn, A. E., Thompson-Schill, S. L., & Bassett, D. S. (2017). Process reveals structure: How a network is traversed mediates expectations about its architecture. Scientific Reports, 7(1), 12733. 10.1038/s41598-017-12876-5

37. Khouri, L., & Nelken, I. (2015). Detecting the unexpected. In Current Opinion in Neurobiology (Vol. 35). 10.1016/j.conb.2015.08.003

38. Lehmann, E. L. (1998). Nonparametrics: Statistical Methods Based on Ranks, Revised. Prentice Hall.

39. Lynn, C. W., Kahn, A. E., Nyema, N., & Bassett, D. S. (2020). Abstract representations of events arise from mental errors in learning and memory. Nature Communications, 11(1), 2313. 10.1038/s41467-020-15146-7

40. Maheu, M., Dehaene, S., & Meyniel, F. (2019). Brain signatures of a multiscale process of sequence learning in humans. ELife, 8, e41541. 10.7554/eLife.41541

41. Maheu, M., Meyniel, F., & Dehaene, S. (2022). Rational arbitration between statistics and rules in human sequence processing. Nature Human Behaviour, 6(8), 1087–1103. 10.1038/s41562-021-01259-6

42. Melloni, L., Schwiedrzik, C. M., Müller, N., Rodriguez, E., & Singer, W. (2011). Behavioral/Systems/Cognitive Expectations Change the Signatures and Timing of Electrophysiological Correlates of Perceptual Awareness. Journal of Neuroscience, 31(4), 1386– 1396. 10.1523/JNEUROSCI.4570-10.2011

43. Meyniel, F., & Dehaene, S. (2017). Brain networks for confidence weighting and hierarchical inference during probabilistic learning. Proceedings of the National Academy of Sciences, 114(19), E3859– E3868. 10.1073/pnas.1615773114

44. Milne, A. E., Wilson, B., & Christiansen, M. H. (2018). Structured sequence learning across sensory modalities in humans and nonhuman primates. Current Opinion in Behavioral Sciences, 21, 39–48. 10.1016/j.cobeha.2017.11.016

45. Milne, A., Zhao, S., Tampakaki, C., Bury, G., & Chait, M. (2021). Sustained pupil responses are modulated by predictability of auditory sequences. Journal of Neuroscience, 41(28), 6116–6127. 10.1523/JNEUROSCI.2879-20.2021

46. Moore, B. C. J., & Gockel, H. E. (2012). Properties of auditory stream formation. In Philosophical Transactions of the Royal Society B: Biological Sciences (Vol. 367, Issue 1591). 10.1098/rstb.2011.0355

47. Ohlenforst, B., Wendt, D., Kramer, S. E., Naylor, G., Zekveld, A. A., & Lunner, T. (2018). Impact of SNR, masker type and noise reduction processing on sentence recognition performance and listening effort as indicated by the pupil dilation response. Hearing Research, 365, 90–99. 10.1016/j.heares.2018.05.003

48. Planton, S., Van Kerkoerle, T., Abbih, L., Maheu, M., Meyniel, F., Sigman, M., Wang, L., Figueira, S., Romano, S., & Dehaene, S. (2021). A theory of memory for binary sequences: Evidence for a mental compression algorithm in humans. PLOS Computational Biology, 17(1), e1008598. 10.1371/journal.pcbi.1008598

49. Press, C., & Yon, D. (2019). Perceptual Prediction: Rapidly Making Sense of a Noisy World. Current Biology, 29(15), R751–R753. 10.1016/j.cub.2019.06.054

50. Romberg, A. R., & Saffran, J. R. (2010). Statistical learning and language acquisition. Wiley Interdisciplinary Reviews: Cognitive Science, 1(6), 906–914. 10.1002/wcs.78

51. Saffran, J. R., Johnson, E. K., Aslin, R. N., & Newport, E. L. (1999). Statistical learning of tone sequences by human infants and adults. Cognition, 70(1), 27–52. 10.1016/S0010-0277(98)00075-4

52. Schapiro, A. C., Rogers, T. T., Cordova, N. I., Turk-Browne, N. B., & Botvinick, M. M. (2013). Neural representations of events arise from temporal community structure. Nature Neuroscience, 16(4), 486–492. 10.1038/nn.3331

53. Sherman, B. E., Graves, K. N., & Turk-Browne, N. B. (2020). The prevalence and importance of statistical learning in human cognition and behavior. Current Opinion in Behavioral Sciences, 32, 15–20. 10.1016/j.cobeha.2020.01.015

54. Skerritt-Davis, B., & Elhilali, M. (2019). A Model for Statistical Regularity Extraction from Dynamic Sounds. Acta Acustica United with Acustica, 105(1), 1–4. 10.3813/AAA.919279

55. Sohoglu, E., & Chait, M. (2016). Detecting and representing predictable structure during auditory scene analysis. ELife, 7(5), e19113. 10.7554/eLife.19113

56. Southwell, R., Baumann, A., Gal, C., Barascud, N., Friston, K., & Chait, M. (2017). Is predictability salient? A study of attentional capture by auditory patterns. Philosophical Transactions of the Royal Society of London B: Biological Sciences, 372(1714), 20160105.

57. Southwell, R., & Chait, M. (2018). Enhanced deviant responses in patterned relative to random sound sequences. Cortex, 109, 92–103. 10.1016/j.cortex.2018.08.032

58. Summerfield, C., Trittschuh, E. H., Monti, J. M., Mesulam, M.-M., & Egner, T. (2008). Neural repetition suppression reflects fulfilled perceptual expectations. Nature Neurscience, 11, 1004–1006. 10.1038/nn.2163

59. Tóth, B., Kristóf, P., Osy, V., Kovács, P., Háden, G. P., Polver, S., Sziller, I., & Winkler, I. (2023). Auditory learning of recurrent tone sequences is present in the newborn’s brain. Neuroimage, 120384. 10.1016/j.neuroimage.2023.120384

60. Warren, R. M., Gardner, D. A., Brubaker, B. S., & Bashford, J. A. (1991). Melodic and Nonmelodic Sequences of Tones: Effects of Duration on Perception. Music Perception: An Interdisciplinary Journal, 8(3), 277–289. 10.2307/40285503

61. Wilson, B., Spierings, M., Ravignani, A., Mueller, J. L., Mintz, T. H., Wijnen, F., van der Kant, A., Smith, K., & Rey, A. (2020). Non-adjacent Dependency Learning in Humans and Other Animals. Topics in Cognitive Science, 12(3), 843–858. 10.1111/TOPS.12381

62. Winkler, I., & Denham, S. L. (2024). The role of auditory source and action representations in segmenting experience into events. Nature Reviews Psychology, 3(4), 223–241. 10.1038/s44159-024-00287-z

63. Winkler, I., Denham, S. L., & Nelken, I. (2009). Modeling the auditory scene: predictive regularity representations and perceptual objects. Trends in Cognitive Sciences, 13(12), 532–540. 10.1016/j.tics.2009.09.003

64. Winkler., I., Paavilainen, P., Alho, K., Reinikainen, K., Sams, M., & Naatanen, R. (1990). The Effect of Small Variation of the Frequent Auditory Stimulus on the Event-Related Brain Potential to the Infrequent Stimulus. Psychophysiology, 27(2), 228–235. 10.1111/J.1469-8986.1990.TB00374.X

65. Yon, D., & Frith, C. D. (2021). Precision and the Bayesian brain. Current Biology, 31(17), R1026–R1032. 10.1016/j.cub.2021.07.044

66. Zekveld, A. A., Koelewijn, T., & Kramer, S. E. (2018). The Pupil Dilation Response to Auditory Stimuli: Current State of Knowledge. Trends in Hearing, 22, 233121651877717. 10.1177/2331216518777174

67. Zhao, S., Skerritt-Davis, B., Elhilali, M., Dick, F., & Chait, M. (2025). Sustained EEG responses to rapidly unfolding stochastic sounds reflect Bayesian inferred reliability tracking. Progress in Neurobiology, 244, 102696. 10.1016/J.PNEUROBIO.2024.102696

